# The nucleus bypasses obstacles by deforming like a drop with surface tension mediated by lamin A/C

**DOI:** 10.1101/2022.03.10.483838

**Authors:** Aditya Katiyar, Jian Zhang, Jyot D. Antani, Yifan Yu, Kelsey L. Scott, Pushkar P. Lele, Cynthia A. Reinhart-King, Nathan J. Sniadecki, Kyle J. Roux, Richard B. Dickinson, Tanmay P. Lele

## Abstract

Migrating cells must deform their stiff cell nucleus to move through pores and fibers in tissue. Lamin A/C is known to hinder cell migration by limiting nuclear deformation and passage through confining channels, but its role in nuclear deformation and passage through fibrous environments is less clear. We studied cell and nuclear migration through discrete, closely spaced, slender obstacles which mimic the mechanical properties of collagen fibers. Nuclei bypassed slender obstacles while preserving their overall morphology by deforming around them with deep local invaginations of little resisting force. The obstacles did not impede the nuclear trajectory or cause a rupture of the nuclear envelope. Nuclei likewise deformed around single collagen fibers in cells migrating in 3D collagen gels. In contrast to its limiting role in nuclear passage through confining channels, lamin A/C facilitated nuclear deformation and passage through fibrous environments; nuclei in lamin-null (*Lmna*^-/-^) cells lost their overall morphology and became entangled on the obstacles. Analogous to surface tension-mediated deformation of a liquid drop, lamin A/C imparts a surface tension on the nucleus that allows nuclear invaginations with little mechanical resistance, preventing nuclear entanglement and allowing nuclear passage through fibrous environments.

## 1. Introduction

Cells migrate through interstitial spaces of fibrous tissue during key physiological processes like wound healing and cancer cell invasion. During this process, the cells have to deform the nucleus to fit through interstitial gaps that are typically smaller than the nuclear size. ^[1–3]^ Deformability of the nucleus has been shown to limit the passage of cells through micropores in the 3D fibrous extracellular matrix. ^[1, 2, 4, 5]^ As such, there is much interest in understanding how the cell can deform the nucleus to fit through interstitial gaps despite its high mechanical stiffness. ^[6–9]^

Nuclear lamins are known to be the key contributors to the mechanical stiffness of the nucleus. ^[10–18]^ Lamin A/C, and not lamin B1, limits the passage of the nucleus through confining pores in 3D collagen gels or microfabricated microchannels. ^[19, 20]^ Furthermore, the depletion of lamin A/C but not lamin B1 significantly softens the nucleus. ^[21]^ These studies collectively suggest that the mechanical stiffness conferred onto the nucleus by lamin A/C impedes nuclear deformation and hence cell passage through narrow micropores or channels.

While the role of lamin A/C in limiting nuclear deformation and passage through confining channels has been studied extensively in pores and microchannels with a smooth contiguous surface, cells such as fibroblasts and cancer cells encounter slender extracellular matrix fibers as they migrate through interstitial tissue. ^[22]^ Here we examined the role of lamin A/C in cell migration through discrete, closely spaced obstacles designed such that their stiffness was similar to the stiffness of single collagen fibers. Unlike in micropores, where the presence of lamin A/C impedes translocation of nuclei, our results show a facilitating role for lamin A/C in the nuclear passage in between slender obstacles. Specifically, wild-type nuclei containing lamin A/C can bypass the slender obstacles while preserving their overall oval shape despite deep local indentations caused by the obstacles. Nuclei lacking lamin A/C, in contrast, become extremely deformed and entangled around obstacles, thus impeding nuclear motion. Our results support a model in which the nucleus deforms like a drop with surface tension conferred by lamin A/C.

## 2. Results

We cultured fibroblasts on an array of fibronectin-coated, flexible, vertical PDMS micropost barriers, which were continuous with the underlying PDMS surface (**schematic in Figure 1A**). The diameter of the microposts (**~1 micron, Figure 1B**) and Young’s modulus were chosen such that their flexural rigidity was similar to that of single collagen fiber bundles (see methods). The microposts were arranged in circular patterns (**Figure 1B**) such that the distance between adjacent pairs of microposts in the pattern was smaller than typical nuclear diameters with a height similar to typical nuclear heights (~5 microns) in cultured cells (**Figure 1B and S1A**). Cells cultured on these fibronectin-coated substrates adhered to the bottom surface and internalized the microposts such that the microposts protruded above the apical cell surface (**SEM image in Figure 1C, yellow arrowheads**).

**Figure 1:**
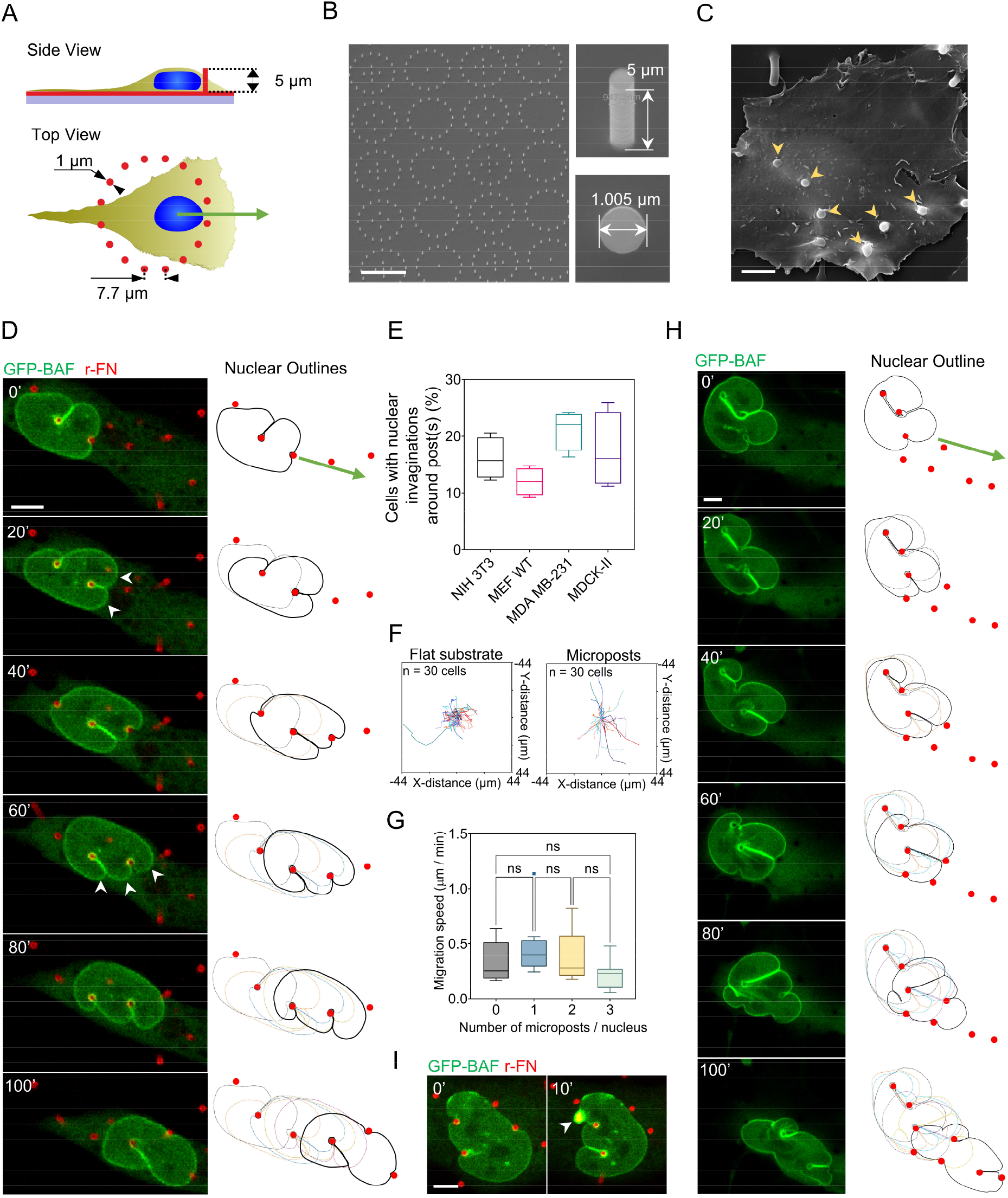
Nuclear invaginations around microposts permit unhindered forward motion of the nucleus. (A) Schematic illustrates micropost geometry relative to cell shape. (B) SEM image shows the circular pattern of fabricated microposts (left, scale bar is 20 μm) and micropost geometry (right). (C) SEM of an MDCK-II cell on microposts, yellow arrowheads indicate internalized microposts (Scale bar is 5 μm). (D) Time-lapse confocal images of an NIH 3T3 fibroblast stably expressing GFP-BAF deforming around 5 μm tall rhodamine-fibronectin coated PDMS microposts (red) and forming transient deep, local invaginations in the nuclear surface (Scale bar is 5 μm). Nuclear outlines relative to the position of microposts are shown on the right. (E) Plot shows the percentage of cells with nuclei that formed deep invaginations around the microposts in different cell types, including fibroblasts (NIH 3T3 and MEF), epithelial cells (MDCK-II), and breast cancer cells (MDA-MB-231); n = 50 cells per condition from 3 independent experiments. (F) Nuclear trajectories of NIH 3T3 fibroblasts migrating on a flat substrate (left) and microposts (right); n = 30 cells imaged for 2 hours for each condition from 3 independent experiments. (G) Bar graph shows the mean nuclear speed in cells migrating against a varying number of microposts. Error bars represent SEM. (ns p>0.05; Brown-Forsythe and Welch ANOVA test). (H) Time-lapse confocal images of an NIH 3T3 fibroblast stably expressing GFP-BAF deforming around 11 μm tall Si microposts (red), forming deep invaginations in the x-y plane, separated by lobes of nearly constant curvature (Scale bar is 5 μm). Nuclear outlines relative to the position of microposts are shown on the right. (I) Nuclear envelope rupture in fibroblasts caused by micropost indentation. White arrowhead points to the local enrichment of GFP-BAF indicative of rupture (Scale bar is 5 μm).

We performed time-lapse confocal fluorescence imaging of fibroblasts stably expressing GFP-BAF (Barrier-to-Auto-Integration Factor) on the micropost substrates. To calculate the force due to cellular or nuclear contact, the microposts were coated with rhodamine-fibronectin (r-FN), and their deflection was tracked by imaging their top and bottom positions. GFP-BAF was imaged as a marker of the nuclear envelope and of envelope rupture. ^[23, 24]^ Despite the fact that the nucleus was wider than the gap in between the micropost obstacles, the nucleus was able to move unimpeded past the obstacles because contact with each new obstacle created a transient deep, local invagination in the nuclear surface (**Figure 1D and Movie 1**). These invaginations continually formed and disappeared as the nucleus encountered microposts during its forward motion. Notably, despite the continual formation and disappearance of the deep invaginations from different directions, the overall oval nuclear shape was preserved (**outlined in Figure 1D**). In many cases, the moving, deforming nuclei slid over the top of the microposts (**Figure S1B, white arrowheads**) without collapsing them (**Figure S1B**) (which gave the appearance that the microposts were passing completely through the nucleus in the lower x-y planes). Even in these cases, the overall oval shape of the nucleus was maintained. Notably, the lobes on either side of the invaginations had near-constant curvature (**Figure 1D, white arrowheads**). This suggests that the lobes are pressurized with the tension in the curved lamina balancing the pressure difference across the nuclear envelope (by Law of Laplace).

Because nuclear deformation is a dynamic phenomenon, only 20-30% of cells that were fixed at a given instant were observed to contain nuclei with deep local invaginations which surrounded the microposts; such invaginations were observed consistently across mesenchymal, epithelial, and cancer cell types (**Figure 1E**). A comparison of cellular trajectories on flat PDMS substrates compared to micropost substrates confirmed that cellular migration was unimpeded on micropost substrates (**Figure 1F**). Nuclear speeds across microposts were 0.34±0.03 μm.min^-1^ and were insensitive to the number of microposts (**Figure 1G**). Speed insensitivity to the number of microposts provides a further indication that invaginations around microposts permit unhindered forward motion of the nucleus.

To determine whether nuclear motion through the micropost array required deflection of microposts and/or moving over the top of the microposts, we repeated these experiments with silicon microposts that were taller but more rigid (11 microns in height) (**Figure 1H and Movie 2**). The nucleus similarly formed deep invaginations in the x-y plane, separated by lobes of nearly constant curvature; these invaginations allowed the nucleus to move unimpeded in between the silicon microposts. Despite the consistent formation of deep nuclear invaginations on the 5-micron tall PDMS microposts and 11-micron tall silicon microposts, rupture of the nuclear envelope, as indicated by local BAF accumulation, was rare (**~5% of cases, Figure 1I**). In contrast, we have previously shown that rapid elastic deformations of the nucleus, even if small, typically cause envelope rupture, ^[25]^ and rupture is more frequent when the nucleus deforms during migration through confining channels. ^[2, 26]^ In the few instances that rupture did occur, the rupture did not typically occur along the invagination (**Figure S1D**). These results with PDMS as well as silicon microposts show that the principal behaviors observed on the microposts (deep nuclear invaginations and unimpeded motion of the nucleus) did not require microposts to be elastically compliant.

All microposts that were internalized by cells were surrounded by the plasma membrane throughout their vertical length (**Figure 2A**). Importantly, the plasma membrane continued to surround microposts even when these microposts were present inside nuclear invaginations. Furthermore, internalized microposts were surrounded by F-actin inside nuclear invaginations (**Figure 2B**). The microposts did not appear to affect the assembly of F-actin networks into basal stress fibers which formed around them (**Figure 2C**). Both outer and inner nuclear membranes surrounded the microposts (**Figure 2D**). The nuclear lamina, visualized by GFP-lamin A expression, behaved similarly to the GFP-BAF labeled nuclear surface around the microposts (**Figure 2E**), with deep invaginations around the microposts (**Figure 2E, yellow arrowheads**). These results show that the microposts are not exposed to cellular or nuclear contents and that the invaginations are able to form even in the stiff nuclear lamina.

**Figure 2:**
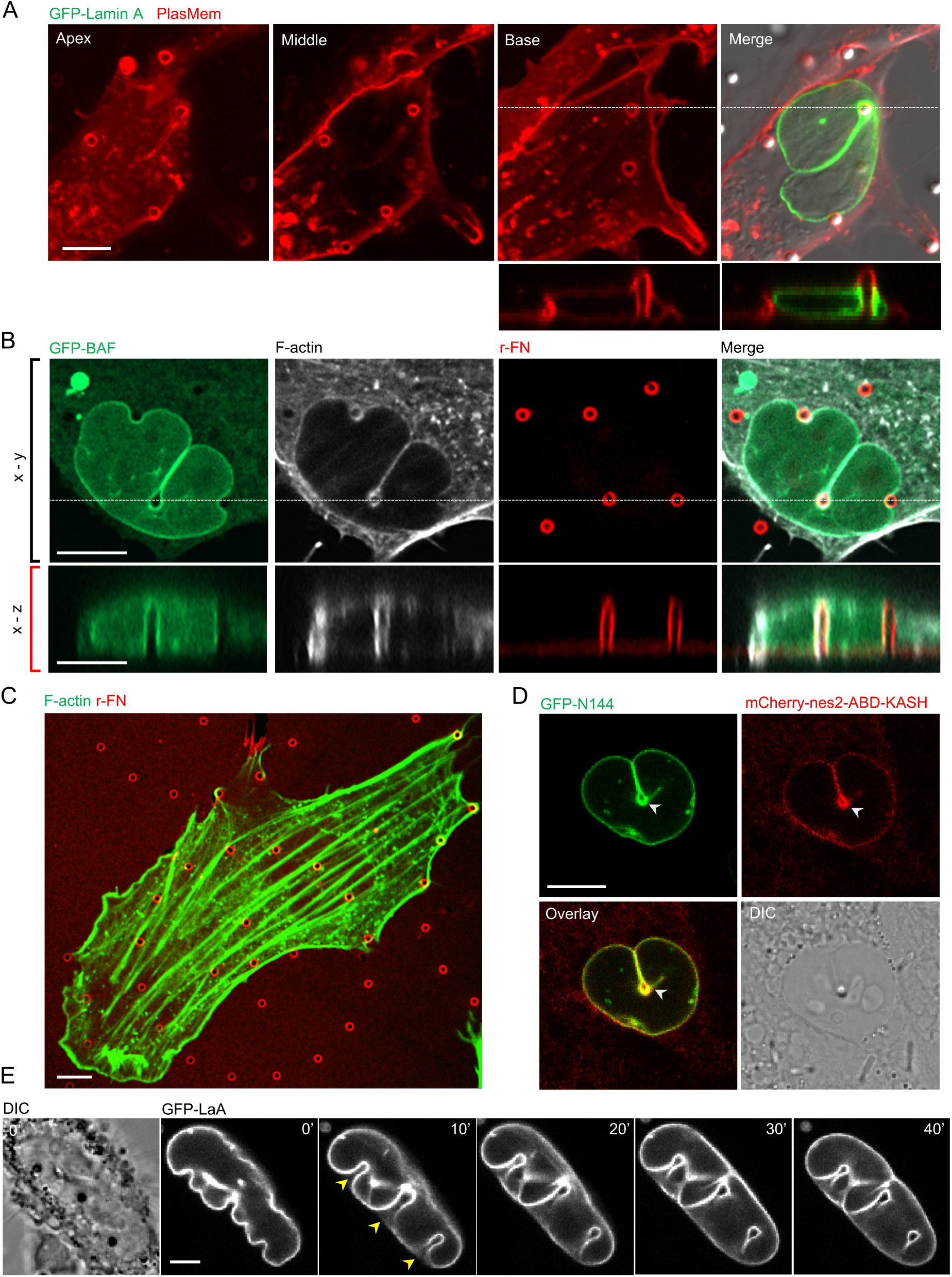
Microposts in nuclear invaginations are not exposed to cellular or nuclear contents. (A) Confocal images of a fibroblast cultured on microposts, with plasma membrane labeled with PlasMem bright red, and expressing GFP-lamin A (green). The apical, middle and basal planes are shown along with the corresponding x-z view situated along the white-dashed line (Scale bar is 5 μm). (B) Confocal images of F-actin stained with phalloidin (white) and GFP-BAF (green) expressing fibroblast with nucleus deformed around the microposts coated with Rhodamine-fibronectin (red); corresponding x-z views are also shown. (Scale bar is 5 μm). (C) Confocal image of F-actin stress fibers (green) in the basal plane of a fibroblast cultured on rhodamine-fibronectin-coated microposts (red) (Scale bar is 5 μm). (D) Images of an MCF-10A cell stably expressing GFP-N144 (green) and mCherry-nes2-KASH (red) cultured on microposts. White arrowheads point to the ring of the inner and outer nuclear membrane proteins around a single micropost present in a nuclear invagination (Scale bar is 5 μm). (E) Time-lapse image sequence showing the development of nuclear invaginations (yellow arrowheads) in a GFP-lamin A expressing fibroblast. (Scale bar is 5 μm).

As the nucleus is generally considered to be mechanically stiff, ^[15, 18, 27–29]^ deep invaginations would be expected to result in a large opposing force that would impede nuclear motion. We, therefore, set out to calculate the magnitude of forces that cause nuclear invagination by measuring the deflection of the microposts with known mechanical stiffness. To quantify micropost deflection in the direction normal to the invaginating nuclear surface, we measured the deflection vector at the top of the micropost and how it was modified following contact with the nuclear surface (**Figure 3A, brown and red arrows**). The differential deflection vector was calculated by vectorial subtraction of the deflection vector at a given time point from the deflection vector before nuclear contact (**blue arrows in Figure 3B**). The component of the differential deflection vector along the direction of the invagination yielded the force exerted by the micropost on the nucleus (**Figure 3A**). The average force was ~2 nN for an indentation depth of 4-6 microns (**Figure 3A–3E**), which is significantly smaller than the corresponding forces measured by AFM (10-15 nN ^[30–32]^). Furthermore, the force appeared to correlate with the invagination depth only up to an initial penetration of ~3 microns, after which it became insensitive to the invagination depth (**Figure 3E**).

**Figure 3:**
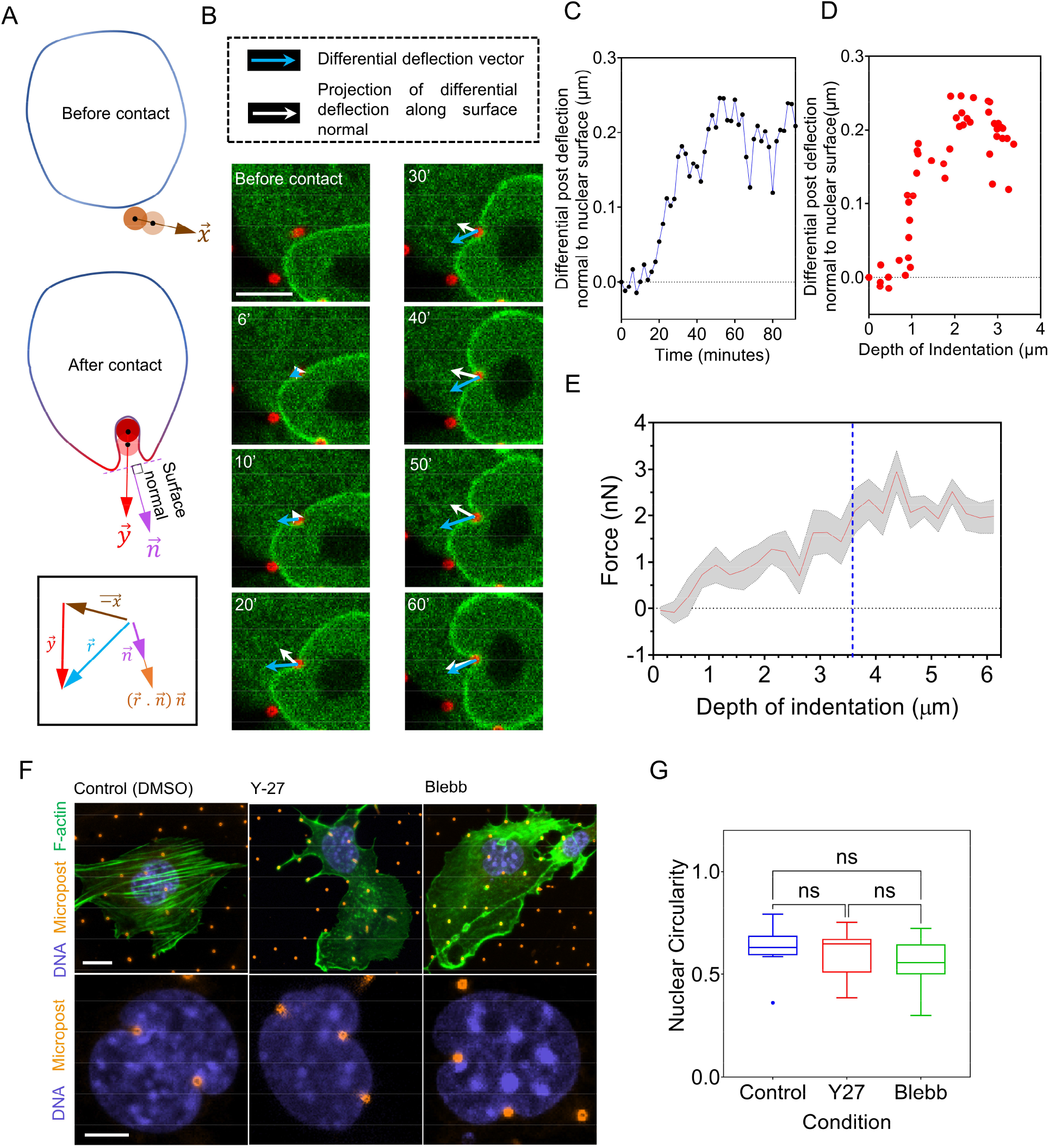
Measurement of forces associated with nuclear invagination and myosin-insensitivity of nuclear invaginations. (A) Schematic shows the calculation of a differential deflection vector 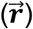 as the vectorial change in deflection of the micropost after nuclear contact, and subsequent projection onto the nuclear surface normal. (B) A representative example of the calculated differential deflection vector (blue) and the corresponding normal component (white arrows) in a nuclear invagination (Scale bar is 5 μm). (C) Plot of the magnitude of the component of the differential vector along the surface normal against time for the example in (B). Time t = 0 refers to the time-point before the first nucleus-micropost contact. (D) Plot of the differential vector along the surface normal corresponding to 3C against the depth of indentation. (E) Plot shows the magnitude of force for mean values pooled from n = 10 cells from at least three independent experiments. Grey area represents SEM. The vertical dashed line (blue) indicates a plateau in the force. (F) Images show representative examples of fibroblasts on microposts (orange) treated with DMSO (control), Y-27632 (25 μM), and Blebbistatin (50 μM) for 12 hours before fixation, followed by staining with Hoechst H33342 (blue, DNA) and Phalloidin (green, F-actin) (Scale is 10 μm in the top panel; Scale bar is 5 μm in bottom panel). (G) Plot of nuclear circularity of control and treated cells in (F), (ns p>0.05; Brown-Forsythe and Welch ANOVA test).

The low magnitude of resisting force to invagination may explain why the invaginations do not hinder the forward motion of the nucleus. Consistent with the low force, the nuclei were able to deform around the posts like the control even upon treatment with Y-27632, a Rho kinase inhibitor, or Blebbistatin, a myosin inhibitor, both of which inhibit actomyosin contractility (**Figure 3F and 3G**). Moreover, the plateau in the force-invagination length curve (**Figure 3E, dashed blue line**) is more consistent with the behavior of an invaginating liquid drop with a surface tension rather than an invaginating elastic solid. The resisting force of a solid is expected to increase with increasing indentation depth, whereas the resisting force of a liquid with constant surface tension is expected to plateau once the invagination is fully developed to the point where two sides of the invagination become parallel. We tested this intuition by indenting the nucleus with the ~ 1-micron tip of a Tungsten microneedle. Rapid deformation of the nucleus in ~15 s with the microneedle produced shapes that resembled kidney-bean-like morphologies (**Figure 4A, top panel**). This response is in contrast to the slower deformations around the microposts over tens of minutes during migration, which produced invaginations closely wrapped around the nucleus (**Figure 4B**). These differences in shape can be explained by a mechanical resistance to a strain of the nuclear interior on the shorter time scale but an overall nuclear pressure balanced by surface tension on the longer time scale. This explanation is consistent with the notion that any mechanical energy stored in the strained nuclear interior dissipates on the time scale of migration such that only the surface tension and the resistance of the nucleus to volume change (i.e., nuclear pressure) govern nuclear deformation in response to external forces on this time scale. ^[7, 33]^

**Figure 4:**
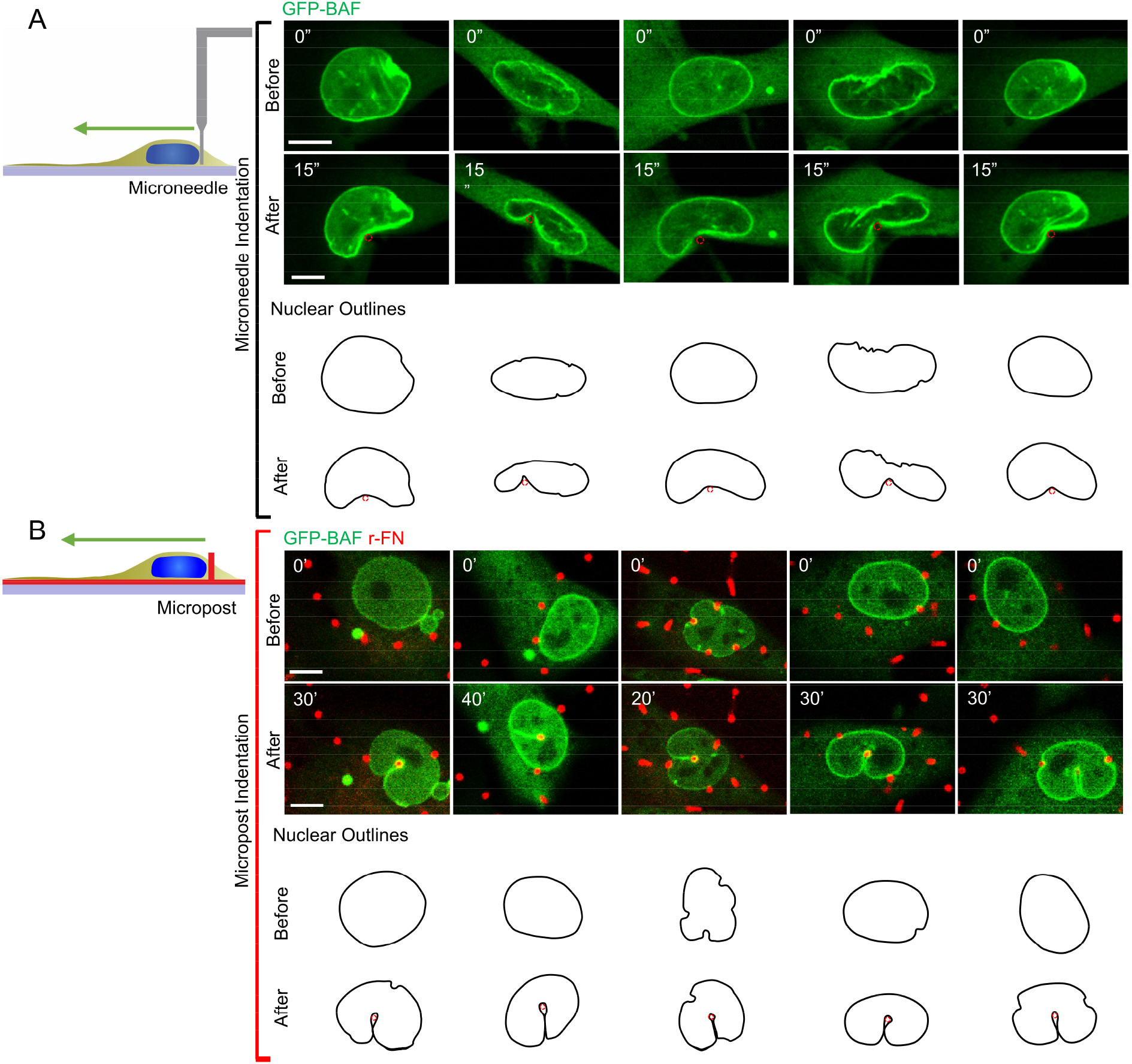
Comparison of nuclear deformation at slow and fast time scales. (A) Images of nuclei of GFP-BAF expressing fibroblasts deformed with a 1-micron diameter Tungsten microneedle at fast time scales (~15 seconds). (B) shows nuclei deformed around microposts in the same cell type over tens of minutes. The nuclear outlines are rotated to highlight the qualitative difference between the nuclear deformation at short (seconds) and long (minutes) time scales (Scale bar is 5 μm).

We explored how these findings relate to three-dimensional cell migration through fibrous environments by examining nuclear deformation in 3-D collagen gels. In these experiments, we observed many instances where a single collagen fiber (~0.4 μm in diameter) caused an invagination in the cell nucleus (**Figure 5A and Movie 3**), much like the microposts (**Figure 5B**). Invaginations were present throughout the thickness of the nucleus (**compare Figure 5C and 5D**). These invaginations allowed the nucleus to bypass the fiber without getting entangled during cell migration (**compare Figure 5E and 5F**). Much like nuclear bypassing of microposts, these results show that local invaginations facilitate nuclear motion around slender obstacles in a 3-D fibrous microenvironment.

**Figure 5:**
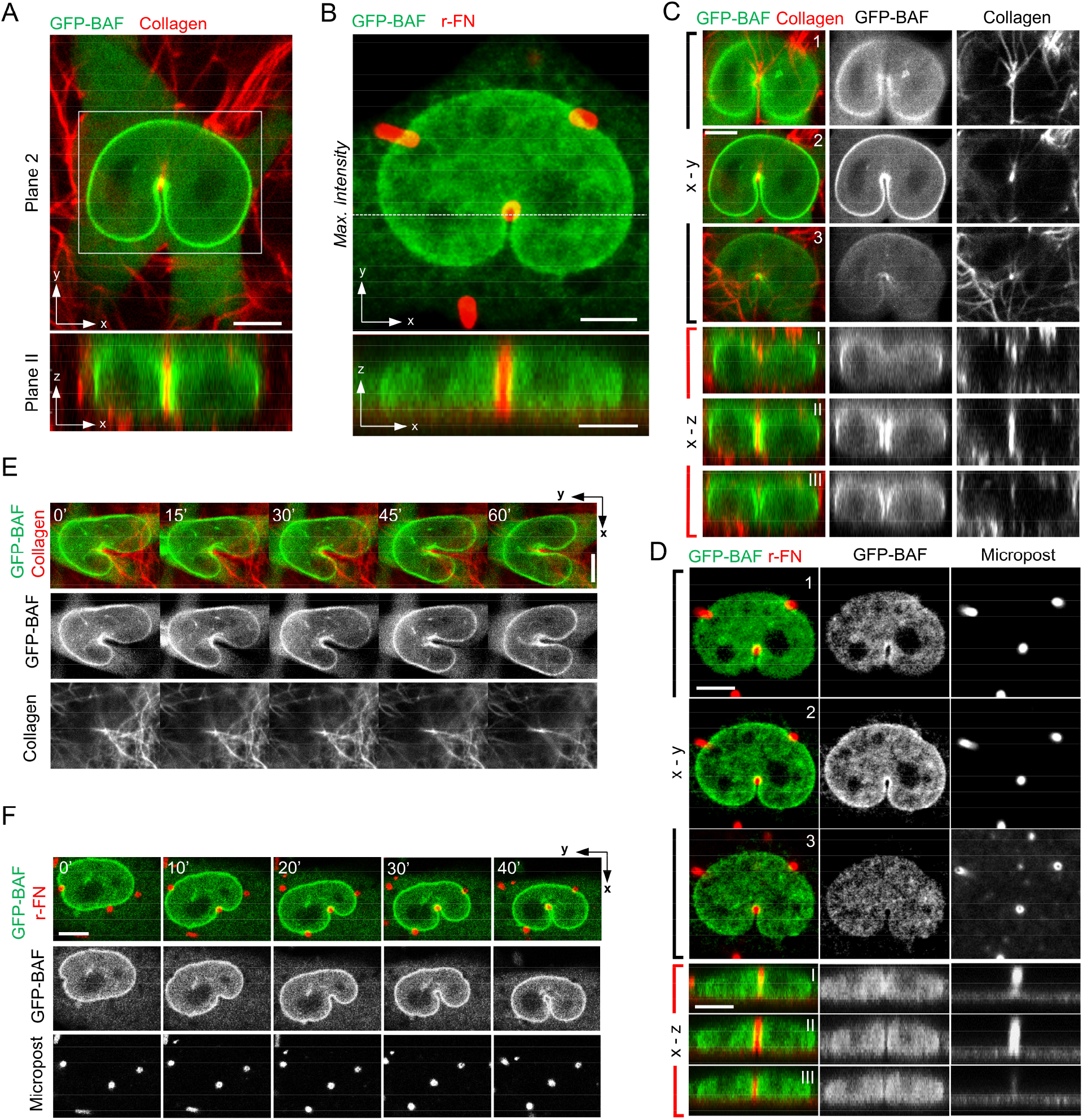
Nuclear invaginations around single collagen fibers during 3D migration. (A) Representative images of a fibroblast (expressing GFP-BAF, green) cultured in 0.5 mg/mL 3-D collagen gel (collagen fibers labeled with NHS ester dye in red). Top (X-Y Plane 2 of example in 5C) and bottom (X-Z Plane II of example in 5C) panels show the horizontal and vertical crosssection views (Scale bar is 5 μm). (B) (Top) Maximum intensity projection of a fibroblast stably expressing GFP-BAF (green) deformed against microposts (red). (Bottom) X-Z reconstruction of confocal z-stacks shows nuclear envelope (green) and microposts (red) in the axial direction (Scale bar is 10 μm). (C) Different cross-sectional views for the nuclear region within the white box in figure 5A are shown in the middle column. X-Y Planes 1-3 show the horizontal crosssection planes focused on the top, middle, and bottom of the nucleus, respectively. X-Z Planes I-III are the vertical cross-section planes located behind, at, and in front of the vertical collagen fiber (Scale bar is 5 μm). (D) Different cross-sectional views of the nucleus are shown in figure 4B. X-Y planes 1-3 correspond to the focal plane of the top, middle, and bottom of the nucleus, respectively (Scale bar is 5 μm). X-Z Planes I-III are the vertical cross-section planes located behind, at, and in front of the vertical micropost (Scale bar is 5 μm). (E) and (F) show time-lapse images of GFP-BAF expressing nuclei deforming around a vertical collagen fiber (E) (Scale bar is 5 μm) or a micropost (F) during cell migration (Scale bar is 10 μm).

As our results suggest that the surface tension and nuclear pressure govern nuclear deformation during encounters with micropost obstacles, we explored the extent to which the nuclear lamina confers these properties on the nucleus. It is possible that the intermediate filament family protein lamin A/C is responsible for surface tension because lamin A/C is a key determinant of the mechanical properties of the nucleus. ^[10–18]^ We examined the shapes of nuclei as they deformed around microposts in migrating mouse embryonic fibroblasts lacking the *Lmna* gene that encodes for lamin A/C (*Lmna*-/- mouse MEFs). In contrast to nuclei in wild type (WT) MEFs, which retained their overall elliptical shape in the x-y plane despite many invaginations wrapped around microposts (**Figure 6A**), nuclei in *Lmna*^-/-^ MEFs lost their overall oval shape while deforming around microposts. Instead, the nuclei had wispy finger-like extensions between microposts that did not wrap around the microposts (**marked with white arrows in Figure 6A, see also Figure S1C**). These behaviors are consistent with diminished surface tension and a corresponding lack of nuclear pressure in *Lmna*^-/-^ MEFs. The extreme nuclear deformation was reflected in a sharply lower nuclear circularity in *Lmna*^-/-^ cells compared to WT cells cultured on microposts (**Figure 6B**). Live cell imaging revealed that the *Lmna*^-/-^ nuclei became entangled around the microposts in a way that prevented their movement past them (**Figure 6C, second panel, Movie 4**). This was in stark contrast to WT nuclei which bypassed the microposts by forming deep local invaginations as before while preserving overall nuclear shape (**Figure 6C, first panel, Movie 5**). The behavior of the WT nucleus could be rescued in *Lmna^-/-^* cells by expressing WT GFP-lamin A/C in them (**Figure 6C, third panel, and Movie 6**). Together, the results suggest that lamin A/C supports a surface tension that balances the nuclear pressure. This surface tension permits the formation of local invaginations without entangling the nucleus, allowing forward nuclear motion while preserving its overall shape.

**Figure 6:**
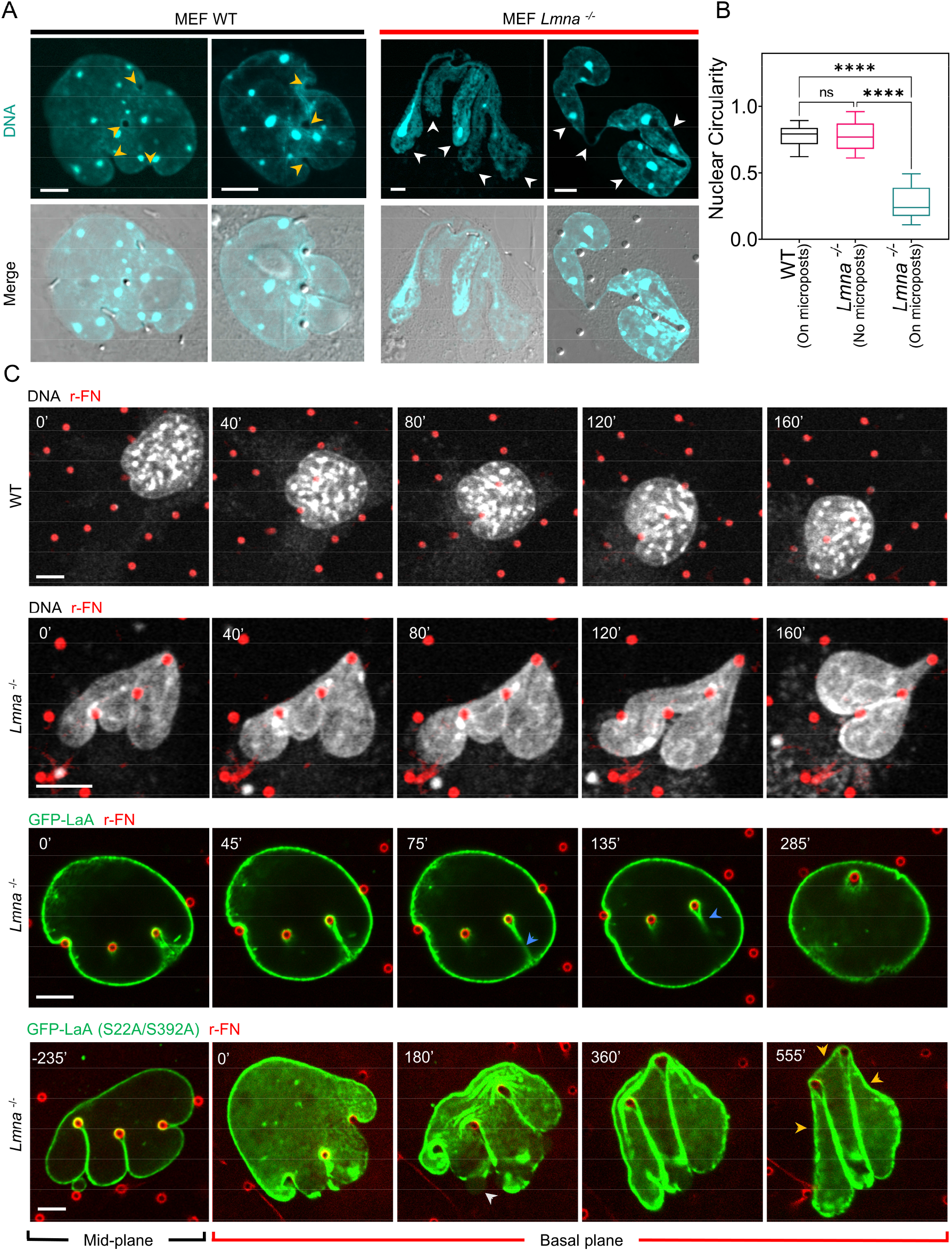
Lamin A/C preserves overall nuclear shape during nuclear invagination around microposts. (A) Images of fixed Hoechst 33342 stained MEF WT and MEF *Lmna^-/-^* nuclei deformed around microposts. Yellow arrowheads point to the micropost locations, and white arrowheads indicate wispy finger-like nuclear extensions around the microposts (Scale bar is 5 μm). (B) Comparison of circularity of MEF WT nuclei and MEF *Lmna^-/-^* nuclei, deformed around microposts, or on surfaces devoid of microposts; (n = 31 nuclei for WT, n = 30 nuclei *Lmna^-/-^* on flat surface, and n = 32 nuclei for *Lmna^-/-^* on microposts from at least three experiments per condition; ns: p > 0.05 and ****: p < 0.0001; Kruskal-Wallis test). (C) Top two panels: time-lapse sequences of a MEF WT nucleus and an MEF *Lmna^-/-^* nucleus stained with NucSpot Live 650 (a live-nuclear imaging dye) (white) during nuclear invagination around rhodamine-labeled microposts (red). Scale bar is 5 μm. Third panel: time-lapse sequence of an MEF *Lmna^-/-^* nucleus expressing WT GFP-Lamin A (green) deforming against a rhodamine-fibronectin labeled micropost (red). Scale bar is 5 μm. Bottom panel: time-lapse sequence of an MEF *Lmna*^-/-^ nucleus expressing GFP-Lamin A (S22A/S392A mutant) (green) deforming against a rhodamine-fibronectin labeled micropost (red). White arrowhead points to a site of blebbing followed by nuclear envelope rupture (rupture is clear in the corresponding movie 9), and yellow arrows point to regions of near-zero and negative curvatures (Scale bar is 5 μm).

*Lmna^-/-^* cells expressing GFP – Lamin A S22A/S392A, a non-phosphorylatable lamin A/C mutant, could initially deform with deep invaginations while preserving the oval nuclear shape around the microposts, similar to the wild-type and unlike the *Lmna^-/-^* nuclei (we confirmed that the mutant lamin A localizes primarily to the lamina (**Figure S2**) consistent with previous reports ^[34]^). This suggests that unlike *Lmna^-/-^* nuclei, Lamin A S22A/S392A expressing nuclei can build up a nuclear pressure and support a surface tension. However, the nuclei were ultimately unable to bypass the microposts, getting entangled in them, like the *Lmna*^-/-^ cells (**Figure 6C, last panel and Movie 7**). These entanglements tended to coincide with frequent rupture events (**Figure 6C and Movie 7, white arrowhead**). Rupture coincided with an apparent transient loss of pressure, as evident from regions of near-zero and negative curvatures (**yellow arrows in Figure 6C**), and caused a loss of the overall oval nuclear shape. These results suggest that nuclear entanglements can also be caused by transient loss of pressure and corresponding loss of surface tension caused by nuclear envelope ruptures.

## 3. Discussion

We observed that the nucleus is able to bypass the microposts by permitting deep local invaginations while maintaining the overall oval nuclear shape; this permitted unhindered forward motion of the nucleus through the micropost array. The regions of near-constant curvature in the lobes on either side of the invaginations suggest a surface tension in the nuclear surface which balances intranuclear pressure. This surface tension and pressure are mediated largely by lamin A/C because *Lmna^-/-^* nuclei deform with long extensions suggestive of a lack of resistance to areal expansion and get entangled on the microposts. *Lmna^-/-^* nuclei thus are never unable to build a pressure owing to a lack of a surface structure that supports a tension to resist the pressure. In contrast to *Lmna^-/-^* nuclei, Lamin A S22A/S392A expressing nuclei are able to initially support a pressure, but eventually lose the pressure due to rupture, resulting in similar entanglements around the microposts as *Lmna^-/-^* nuclei.

The motion of the nucleus through the micropost array (**Movie 1**) is reminiscent of the behavior of a liquid drop encountering slender obstacles (**Movie 8**). This liquid-drop like behavior is demonstrated more directly in **Figure 7, top panel, and Movie 8**, which shows an oil drop with higher surface tension deforming with a narrow, local invagination around the object such that apposing surfaces come together in the wake of the object while the overall oval shape of the drop is preserved. In contrast, the indentation of the oil drop with a lower surface tension causes a more extreme shape distortion from the starting oval shape (**Figure 7, bottom panel, and Movie 9)**. In an analogous manner, lamin A/C appears to create a nuclear surface tension that allows bypassing of the nucleus around slender objects by permitting deep invaginations while maintaining the original overall nuclear shape. Removal of lamin A/C appears to reduce the surface tension, as evident from the distortion of the overall nuclear shape and nuclear entanglement around the microposts.

**Figure 7:**
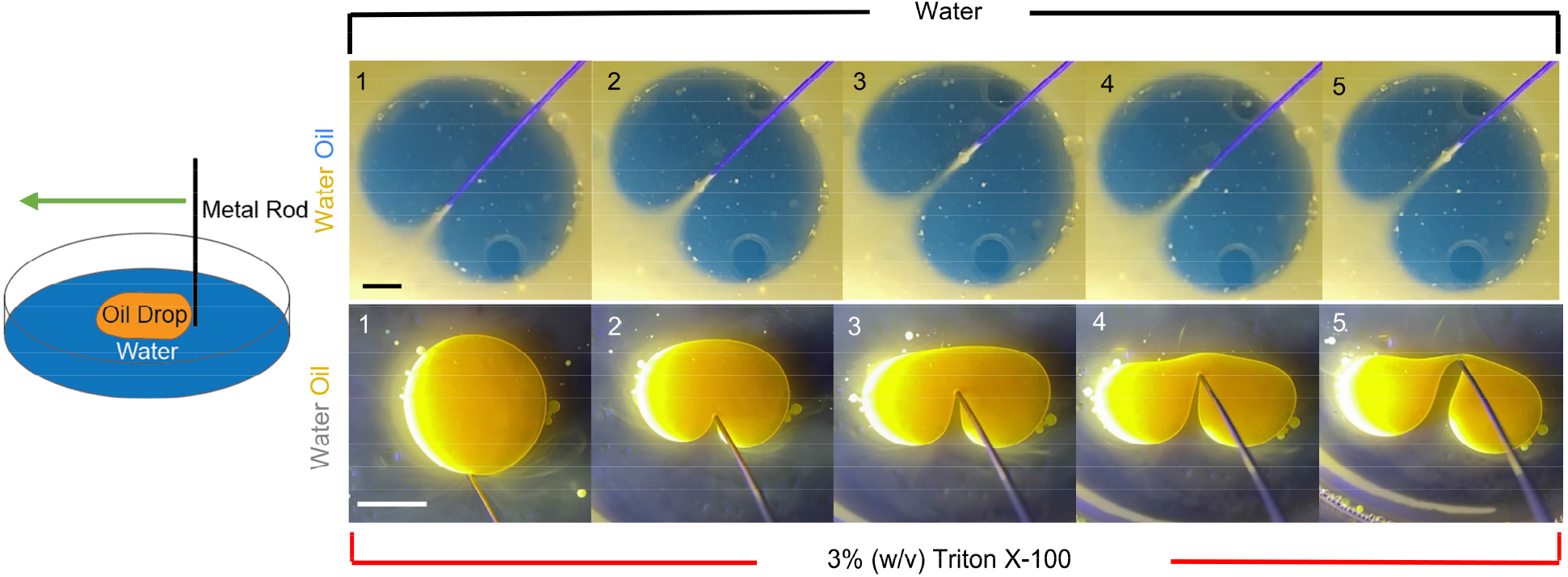
Deformation of an oil drop with a metal wire. (Top) An oil drop (blue) in water (yellow) indented with a metal wire (diameter is 0.5 mm). (Bottom) An oil drop (yellow) in 3% (w/v) Triton X-100 in water indented with the same metal wire (Scale bar is 5 mm).

Unlike an oil drop in water, where the surface tension arises from intermolecular attraction within each phase, surface tension in the nucleus arises due to resistance of the nuclear lamina to areal expansion. The source of tension in the lamina is likely the increase in nuclear pressure upon nuclear flattening in spread cells ^[7, 33]^ as well as the compression of the nucleus against the microposts. Lamin A/C is required to sustain this tension. These results are consistent with our previous finding that lamin A/C limits the flattening of nuclei in spreading cells due to resistance to its areal expansion. ^[35]^

It is well known that on short time scales, nuclei can behave visco-elastically. ^[12, 15, 18, 27, 36–41]^ These measurements are typically on the time scale of several seconds, while our observations are on the order of minutes to hours. Thus, it is difficult to extrapolate the shortterm measurements to the long-timescale behavior. Also, we have previously shown that on the time scale of cell migration, nuclear deformations that occur during migration are not elastic and are primarily limited by the resistance of the lamina to areal expansion and the nuclear volume to compression (^[35, 42]^, and reviewed in ^[7, 33]^). That is, on longer time scales, deformed nuclear shapes in cells are not restored following the removal of cellular forces because of any elastic energy stored in the nuclear shape. However, a pressurized nucleus will tend to maintain an oval shape due to its surface tension, much like a liquid drop.

Our finding of lower forces for more significant deformations of the nuclear surface as compared to AFM measurements taken on a shorter time-scale (seconds), the lack of correlation between force and invagination length at deeper invaginations, the liquid-like behavior of the nucleus where the shape of the lobes between the microposts is determined primarily by the nuclear pressure and surface tension, and the qualitative differences between nuclear shapes indented with a probe in several seconds and nuclear shapes that invaginate over several minutes around microposts in migrating cells, are all also consistent with the notion that any elastic energy stored in the deformed nuclear shape dissipates on the much longer time scale of cell migration. ^[42]^

The three-dimensional extracellular matrix through which cells migrate offers distinct types of barriers to cell migration, including pores and slender fibers. Lamin A/C hinders migration through narrow pores in 3D gels ^[1, 2, 4, 5]^ and confining microfabricated channels. ^[1, 16, 20]^ This is because the tensed lamina resists the areal expansion required for the nucleus to pass through narrow pores and channels. ^[7, 26]^ However, in the context of fibrous environments, our results point to a new mechanism in which nuclear surface tension imparted by lamin A/C facilitates cell migration around slender obstacles by preventing entanglement of the nucleus.

## 4. Materials and Methods

### 4.1 Micropost Design and Fabrication

The height of the microposts was selected such that the micropost height is similar to the nuclear height. ^[43]^ The micropost patterns were designed in AutoCAD 2018. Circles of diameter 1 μm were placed on corners of an equilateral hexadecagon (side length = 7.71 μm). The side length of the hexadecagon was selected such that the area enclosed by the hexadecagon equaled the average cell spreading area (~1200 μm^2^ for NIH 3T3 fibroblasts). The center-to-center distance between the micropost was selected such that the minimum edge-to-edge distance between any two adjacent microposts (6.6 μm) was less than the average diameter of the nucleus (~10 μm for NIH 3T3 fibroblasts). This hexadecagon pattern was arrayed onto an area of 1 cm by 1 cm. A P-type silicon wafer of diameter 100 mm was coated with a 1-micron AZ1512 photoresist. The above AutoCAD design was patterned on a chrome-coated glass mask using a Heidelberg MLA-150 DWL 66 laser writer. The glass mask and wafer were then mounted and exposed on an MA6 contact aligner and then developed using a 4:1 AZ-300MIF solution. Wafer was etched on Oxford DRIE, and the photoresist was removed using O_2_ plasma. Post fabrication, the wafer was coated with (tridecafluoro-1, 1, 2, 2-tetrahydrooctyl)-1-trichlorosilane (T2492, UCT Inc., Bristol, PA, USA) (~50 nm thick using Chemical Vapor Deposition) to ensure low adhesion of PDMS on the wafer and hence promote easy peeling.

A negative mold was prepared from a patterned master silicon wafer using PDMS (Sylgard 184, Dow Corning, Midland, MI, USA) mixed at the base to cross-linking agent ratio of 10:1 (w/w) and cured at 60°C for 2 hours. The negative was peeled from the wafer and cut into blocks. These blocks contained the patterned area. A block was then covered with an evenly spread layer of uncured PDMS (mixed at the base to cross-linking agent ratio of 10:1 (w/w)) and pressed against a No.1.5 glass-bottom dish (FD35-100, WPI Inc., Sarasota County, FL, USA). The dish with the negative mold was cured at 60°C for 2 hours to form the micropost topology on the glass surface.

### 4.2 Flexural Rigidity

Flexural rigidity of collagen fibers or PDMS microposts was calculated as *El* where *E* is Young’s modulus, 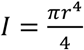 is the second moment of inertia for a cylindrical geometry, and *r* is the radius of the cylinder. *E* for collagen fibrils is reported to be in the range of 100 MPa to 360 MPa ^[44]^. For average fiber diameters of 0.41 ± 0.09 μm in our collagen gels, *EI* = 1.37 × 10^-16^ Nm^2^ to 4.94 × 10^-16^ Nm^2^ for collagen fibers which is roughly comparable to *EI* = 1.11 × 10^-16^ Nm^2^ for PDMS microposts (PDMS mixed at 10:1 base to cross-linker ratio and cured at 60°C for 2 hours) of 1 micron in diameter. Young’s modulus of PDMS was measured experimentally. Samples were prepared as per ASTM D412-C standards. PDMS was mixed at a 10:1 base-to-cross-linking agent ratio and cured at 60°C for 2 hours. Ten samples were tensile tested on an Instron 6800 series universal testing system, and stress-strain curves were acquired and analyzed to determine Young’s modulus of PDMS. The Young’s modulus was measured to be 2.27±0.04 MPa.

### 4.3 Stable Cell Lines and Plasmids

To construct the mCherry-ABD-KASH2-IRES-GFPx3-N144 pBabe puro plasmid, GFPx3-N144 was amplified from GFPx3-N144 pcDNA3.1 ^[45]^ using primers (5’) TGTGGTGGTACGTAGGAATTCGGTTTAAACGCCACCATGGTGAGCAAG (3’) and (5’) ACACACATTCCACAGGGTCGACTTAAGGGGATTC (3’) and inserted into a SnaBI, and SalI cut pBabe puro plasmid. An IRES amplified from pMSCV-IRES-mCherry (a gift from Dario Vignali, Addgene plasmid # 52114) using primers (5’) TGTGGTGGTACGTAGGAATTCGAATTCGCGGGATCAATTCCG (3’) and (5’) CTTGCTCACCATGGTGGCGTTTAAACTTATCGTGTTTTTC (3’) and inserted on the N terminus of GFPx3-N144 pBabe puro. mCherry-ABD-KASH2 pBabe puro was cloned by fusing the actin-binding domain (aa 1-286) to the KASH domain (6825-6884) of human Nesprin 2 (NP_055995.4). This was amplified using primers (5’) TCCGGACTCAGATCTCGAGGCGCATCTAGTCCTGAGCTT (3’) and (5’) TAACTGACACACATTCCACAGGGTCGACCTATGTGGGGGGTGGCCCATTG (3’) and inserted into XhoI, and SalI cut mCherry-NLS pBabe puro plasmid. mCherry-ABD-KASH2 pBabe puro was then amplified with primers (5’) GATCCCAGTGTGGTGGTACGTAGCCACCATGGTGAGCAAGGGCGAGGAGG (3’) and (5’) GGGCGGAATTGATCCCGCGAATTCCTATGTGGGGGGTGGCCCATTG (3’) and inserted into SnaBI, and EcoRI cut IRES-GFPx3-N144 pBabe puro to create mCherry-ABD-KASH2-IRES-GFPx3-N144 pBabe puro. Human lamin A, in frame with GFP, was subcloned into pBabe.puro cut with EcoRI and PmeI using a forward primer TGTGGTGGTACGTAGGAATTCGCCACCATGGTGAGCAAG and a reverse primer CGACTCAGCGGTTTAAACCTACATGATGCTGCAGTT. NIH 3T3 cells stably expressing GFP-BAF WT were created as described previously. ^[45]^ MCF10A breast epithelial cells stably coexpressing GFP-N144 ^[45]^ and mCherry-KASH-ABD-Nes2 and NIH 3T3 cells stably expressing GFP-Lamin A or GFP-53BP1 were generated by retroviral transduction as described. ^[23]^ A-type lamin–deficient (*Lmna*^-/-^) MEFs and WT MEFs were generously provided by Yixian Zheng (Carnegie Institution for Science, Washington, DC, USA) and were derived from *Lmna*-knockout mice ^[46]^ using methods described previously. ^[47]^ GFP-Lamin-A S22A/S392A (a generous gift of the Goldman lab) was described previously. ^[34]^ Lipofectamine 3000 (L3000001, Invitrogen, Carlsbad, CA, USA) was used for transient transfections as per the manufacturer’s recommended protocol.

### 4.4 Cell Culture and Drug Treatment

All cells were maintained in a humidified incubator at 37°C and 5% CO_2_. NIH 3T3 fibroblasts (CRL-1658, ATCC, Manassas, VI, USA) were cultured in Dulbecco’s Modified Eagle’s Medium with 4.5 g.L^-1^ glucose (10-013-CV, Corning, Corning, NY), supplemented with 10% v/v donor bovine serum (16030074, Gibco, Waltham, MA, USA) and 1% v/v penicillin/streptomycin (30-002-CI, Corning, Corning, NY, USA). MCF10A human breast epithelial cells (CRL-10317, ATCC, Manassas, VI, USA) were maintained in DMEM/F12 medium (11039-021, Invitrogen, Carlsbad, CA, USA) supplemented with 20 ng.ml^-1^ epidermal growth factor (AF-100-122, Peprotech, Rocky Hill, NJ, USA), 0.5 mg.ml^-1^ hydrocortisone (50-23-7, Sigma-Aldrich, St. Louis, MO, USA), 100 ng.ml^-1^ cholera toxin (9012-63-9, Sigma-Aldrich, St. Louis, MO, USA), 100 mg.ml^-1^ insulin (11070-73-8, Sigma-Aldrich, St. Louis, MO, USA), 1% v/v penicillin-streptomycin (30-002-CI, Corning, Corning, NY, USA), and 5% v/v horse serum (16050-122, Invitrogen, Carlsbad, CA, USA). MDA-MB-231 (HTB-26, ATCC, Manassas, VI, USA) were cultured in Leibovitz’s L-15 Medium (10-045-CV, Corning, Corning, NY, USA), supplemented with 10% v/v donor bovine serum (16030074, Gibco, Waltham, MA, USA) and 1% v/v penicillin/streptomycin (30-002-CI, Corning, Corning, NY, USA). MDCK (NBL-2) (CCL-3, ATCC, Manassas, VI, USA) cells were cultured in DMEM with 4.5 g.l^-1^ glucose (10-013-CV, Corning, Corning, NY), supplemented with 10% v/v donor bovine serum (16030074, Gibco, Waltham, MA, USA) and 1% v/v penicillin/streptomycin (30-002-CI, Corning, Corning, NY, USA). MEF WT and MEF *Lmna^-/-^* cells were cultured in DMEM with 4.5 g.l^-1^ glucose (10-013-CV, Corning, Corning, NY), supplemented with 15% v/v donor bovine serum (16030074, Gibco, Waltham, MA, USA) and 1% v/v penicillin/streptomycin (30-002-CI, Corning, Corning, NY, USA).

For actomyosin inhibition experiments, cells were seeded on rhodamine-conjugated fibronectin (FNR01, Cytoskeleton Inc., Denver, CO, USA) coated microposts, followed by a replacement of media containing DMSO (control) (BP231-4, Fisher Scientific, Hampton, NH, USA) or 25 μM Y-27632 (Y0503, Sigma-Aldrich, St. Louis, MO, USA) or 50 μM Blebbistatin (B0560, Sigma-Aldrich, St. Louis, MO, USA) within 2 hours after seeding cells. Samples were incubated in the treatment media for 12 hours, fixed with 4% paraformaldehyde (J61899, Alfa Aesar, Haverhill, MA, USA) at room temperature for 15 minutes, and washed thrice with 1X PBS (21-040-CM, Corning, Corning, NY, USA).

### 4.5 Fluorescent Labeling

The micropost pattern was coated with rhodamine-conjugated fibronectin (FNR01, Cytoskeleton Inc., Denver, CO, USA) at a concentration of 5 μg.ml^-1^ to promote cell adhesion while fluorescently-labeling the microposts. For imaging F-actin, plasma membrane, or nuclei in some experiments, cells were fixed in 4% paraformaldehyde (J61899, Alfa Aesar, Haverhill, MA, USA) at room temperature for 15 minutes and washed thrice with 1X PBS (21-040-CM, Corning, Corning, NY, USA). Hoechst (875756-97-1, Sigma-Aldrich, St. Louis, MO, USA) was used to stain DNA, and Alexa Fluor-488 phalloidin (A12379, ThermoFisher Scientific, Waltham, MA, USA) was used to stain F-actin in fixed samples. NucSpot Live 650 (40082, Biotium, San Francisco, CA, USA) was used to stain DNA in live cells. PlasMem Bright Red (P505, Dojindo Molecular Technologies, Inc., Rockville, MD, USA) was used to stain the plasma membrane. All reagents were used at the concentration recommended by the manufacturer.

### 4.6 Microscopy

Imaging was performed on a Nikon Ti2 eclipse laser scanning A1 confocal microscope (Nikon, Melville, NY, USA) with DU4 detector using Nikon CFI Plan Apo Lambda 60X/1.4 NA oil immersion objective lens (MRD01605, Nikon, Melville, NY, USA). Immersion oil Type 37 (16237, Cargille Labs, Cedar Grove, NJ, USA) was used at 37°C (*R.I*. = 1.5238 for *Λ* = 486.1 nm). Alternatively, imaging was performed on an Olympus FV3000 (Olympus Scientific Solutions Americas Corp., Waltham, MA, USA) using Super Apochromat 60x silicone oil immersion lens (UPLSAPO60XS2, Olympus Scientific Solutions Americas Corp., Waltham, MA, USA) or on a Zeiss LSM 900 (Carl Zeiss Jena GmbH, Jena, Germany) with Airyscan 2 using W Plan-Apochromat 20x/1.0 objective (421452-9681-000, Carl Zeiss Jena GmbH, Jena, Germany) or C Plan-Apochromat 63x/1.4 Oil objective (421782-9900-000, Carl Zeiss Jena GmbH, Jena, Germany). For 3D confocal imaging of fixed samples, a pinhole opening of 1 *Airy disk* was selected and a z-step size of 100 nm or 250 nm to ensure overlapping z-stacks while sampling at less than half of the depth of focus (which corresponds to an optical section of ~500 nm for 488 nm light) to satisfy Nyquist criterion and minimize photobleaching artifacts [48]. Live timelapse cell imaging was performed in a heated and humidified chamber (Tokia Hit USA Inc., Montgomery, PA, USA), and cells were maintained at 37°C and 5% CO_2_. For live-cell time-lapse 3D confocal imaging, a pinhole opening of 1.2 *Airy disks* (which corresponds to an optical section of ~600 nm for 488 nm light) and a z-step size of 1 micron was used to minimize phototoxicity and photobleaching of fluorescent probes.

For SEM imaging, PDMS microposts were fabricated on a 2 cm by 2 cm piece of a silicon wafer. Capsules containing PDMS micropost were then dehydrated in a graded ethanol series 25%-100%, at 10-min intervals for each 5% increment, followed by two exchanges of anhydrous ethanol. The dehydrated micropost was loaded into a critical point dryer, Tousimis Autosamdri-815 (Rockville, MD, USA), with CO_2_ as the transition fluid and in stasis mode overnight. The critical-point-dried PDMS microposts were mounted on carbon adhesive tabs on aluminum specimen mounts and rendered conductive with Au/Pd using a DeskV Sputter Coater (Denton Vacuum, Moorestown, NJ, USA). The microposts were imaged using a Hitachi SU5000 Schottky field emission SEM (Hitachi High-Technologies, Schaumburg, IL, USA) operated at 5kV.

### 4.7 Collagen Gels

Gels were designed for low collagen fiber density (0.5 μg.ml^-1^) to increase the probability of constricted migration and large fiber diameters (0.409 ± 0.093 μm) to increase aggregate fiber stiffness. Rat tail type-1 collagen solution of concentration 0.5 μg.ml^-1^ in a complete cell culture medium was neutralized to *pH* = 7.0 with 1N NaOH and pipetted to glass-bottom microplates (MatTek Corporation, Ashland, MA, USA). The collagen solution was allowed to polymerize at 4 °C overnight and then at 37 °C for 30 min to finish the polymerization. The polymerized collagen gel was then labeled with 50 μg.ml^-1^ Alexa Fluor 594-conjugated-NHS ester dye (A20004, Thermo Fisher, Waltham, MA, USA) in *pH* = 8.7 NaHCO3 buffer for 1 h at 37 °C, followed by 3 rounds of 1X PBS wash before seeding the cells. NIH 3T3s expressing GFP-BAF were then seeded onto the labeled collagen gel in a complete cell culture medium and incubated at 37° C overnight before imaging. GM6001 (364206-1MG, Sigma-Aldrich, St. Louis, MO, USA), a broadspectrum matrix metalloproteinase inhibitor, was added to the cell culture medium at a concentration of 10-20 μM to inhibit matrix metallopeptidase activity and minimize collagen fiber degradation. Static or time-lapse confocal images were taken with a Zeiss LSM800 confocal microscope, equipped with an environmental control chamber and a 40X water-immersion lens (*N.A*. = 1.1).

### 4.8 Microneedle Indentation

Nuclei of cells adherent on a glass-bottom dish were deformed using a Tungsten microneedle (MN005S, MicroProbes, Gaithersburg, MD, USA) with a 1-micron tip diameter, bent such that the tip approached nearly perpendicular to the base of the dish. The microneedle was attached to an Injectman 4 micromanipulator (5192000027, Eppendorf, Enfield, CT, USA) to control nuclear deformation.

### 4.9 Oil Drop Experiments

Canola oil was mixed with highlighter ink (in trace amounts) and suspended in water or water with 3% (w/v) Triton X-100 Detergent Solution (85111, ThermoFisher Scientific, Waltham, MA, USA). The oil drop was then visualized in blacklight as it was deformed by a thin steel wire 0.5 mm in diameter.

### 4.10 Nuclear or Micropost Height Measurements

X-Z projections were reconstructed using NIS-Elements AR 5.02.01, and the maximum intensity projection was applied to the reconstructed images. Intensity profiles along the object’s axial direction (in the z-direction) were exported to Origin PRO (OriginLab Corporation, Northampton, MA, USA). A Gaussian non-linear fit based on the Levenberg-Marquardt algorithm was applied to the intensity values, and the top and bottom edges of the nucleus or the micro were determined with the full width at half maximum method (FWHM) ^[49]^ function in Origin. The distance between the top and bottom edge of the nucleus or the micropost was reported as the corresponding height.

### 4.11 Deflection Measurements

We measured how the deflection vector (the vector joining the centroids of the bottom of the micropost with the top of the micropost) of the micropost was modified after contact with the nuclear surface (**Figure 3A, brown and red arrows**). Z-planes corresponding to the top and bottom of the fluorescently labeled microposts were acquired. The deflection of a micropost was calculated with an automated program written in MATLAB 2019a (MathWorks, Natick, MA, USA). Centroid-detection-based particle-tracking routines described earlier ^[50]^ were employed to detect and quantify the micropost-coordinates. Briefly, a bandpass filter and a brightness threshold were applied to the raw images to isolate the micropost from the background. The centroid position of each micropost was calculated by fitting 2-D Gaussians to the pixel-intensities around the object of interest, where the peak position of the Gaussian fit was reported as the centroid. Vectorial subtraction between the centroid of the bottom of the micropost and the top of the micropost was reported as a deflection vector. Differential deflection vector was calculated by vectorial subtraction of the micropost deflection vector just before nuclear contact from the micropost deflection vector at a given time point (**blue arrows in Figure 3B**). For representing the deflection vector on images, arrows were scaled ten times the actual vector. The differential force component normal to the nuclear surface (**Figure 3A, orange arrow**) is the force associated with the local indentation of the nuclear surface due to the micropost. To calculate the differential deflection vector component, a line was drawn to join the two protruding nuclear surfaces around the micropost. This line was then rotated 90° to get the direction normal to the nuclear pocket around the micropost (**see Figure 3A, magenta arrow**). The differential deflection vector component along the direction normal to the nuclear surface in subsequent time frames was calculated and reported as the differential micropost deflection normal to the nuclear surface (**Figure 3B, white arrows**). The magnitude of force was calculated under the assumption of a uniformly distributed force across the length of the micropost, which is reasonable given that the nuclear contact with the micropost is relatively uniform throughout the micropost height (**see Figure S1A**). The uniformly distributed load was calculated as 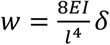 where *δ* is the measured deflection of the top of the micropost relative to micropost bottom, *E* = 2.21 ± 0.04 MPa is Young’s modulus of PDMS mixed at the base to cross-linking agent ratio of 10:1 (w/w) and cured at 60°C for 2 hours, 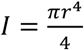 is the area moment of inertia of the circular cross-section of the micropost of radius *r*, and *l* is the length of the micropost. The force associated with a contact by the nucleus was determined by multiplying the uniformly distributed load force per unit length by the length of the microposts (5 μm) and reported as the total force in **Figure 3E**.

### 4.12 Nuclear Speeds

For automated analysis of nuclear speeds, a custom program was written in MATLAB 2019a. First, a threshold was applied to the time-lapse images of nuclei stably expressing GFP-BAF. Then, the nuclei boundaries were traced using MATLAB protocols to filter the image using a feature size-based bandpass filter, binarize (black/white) the image, fill interior gaps, and perform edge-detection. Area and perimeter filters were employed to identify nuclei and differentiate them from noise. Finally, the coordinates of each nuclear centroid were calculated as the mean of X and Y coordinates of the identified nuclear boundary. The centroid displacement against time was fitted with a straight line, and the average of the slopes was reported as the average nuclear speed (**Figure 1G**).

### 4.13 Circularity and Curvature Measurements

The Auto-threshold function in ImageJ was used to determine the nuclear boundaries from images of nuclei fluorescently labeled with Hoechst or GFP-BAF. After thresholding, the Fill Holes function was used to fill holes inside the nuclear bounds and binarize the image. The values of perimeter and area were acquired with the Measure function in ImageJ. Circularity was calculated using the formula, 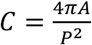 where, *C* is the calculated circularity, *A* is the measured area, and *P* is the measured perimeter. Curvature was measured using the Kappa Curvature analysis plug-in in FIJI. To measure the total surface area of the nucleus, first, confocal z-stacks of Hoechst-stained nuclei were acquired using a z-step size of 250 nm.

### 4.14 Statistical Analysis

GraphPad Prism 9.0 was used for statistical analysis and graphical representations of data. On box plots, the central mark indicates the median, bottom and top edges of the box indicate the 25th and 75th percentiles, respectively, and whiskers extend to the most extreme data point. Statistical tests included Student’s t-test, Brown-Forsythe and Welch analysis of variance (ANOVA) test, Kruskal-Wallis test and Mann-Whitney test. Differences between values were considered statistically significant when p < 0.05 and non-significant (ns) for p > 0.05. The sample size (n) for each statistical analysis is indicated in individual figure legend and the sample sizes used were sufficient to perform statistical analyses. All cell-culture experiments were performed independently at least thrice.

## Movies

Movie 1: **Encounters of a GFP-BAF expressing cell with 5 μm PDMS microposts.** Timelapse movie shows the nucleus of a GFP-BAF expressing fibroblast (green) moved unimpeded past the rhodamine-fibronectin labeled microposts (red) as contact with each new obstacle created a transient deep, local invagination in the nuclear surface (Scale bar is 5 μm).

Movie 2: **Encounters of a GFP-BAF expressing cell with 11 μm Si microposts.** Time-lapse movie shows deep invaginations developing in the nucleus of a GFP-BAF expressing fibroblast (green); the invaginations were separated by lobes of nearly constant curvature that allowed the nucleus to move unimpeded past the microposts (Scale bar is 10 μm).

Movie 3: **Examples of nuclei deforming around single collagen fibers in collagen gels.**

Time-lapse movies of nuclei of GFP-BAF expressing fibroblasts deforming and wrapping around single collagen fibers in 3-D collagen gels or squeezing in-between the interstitial spaces between fibrils (Scale bar is 5 μm).

Movie 4: **Entanglement of *Lmna-/-* nuclei around microposts.** Time-lapse movie of MEF *Lmna^-/^-* nuclei stained with NucSpot Live 650 (a live-nuclear imaging dye, white) as they deform against rhodamine-labeled microposts (red) (Scale bar is 5 μm).

Movie 5: **WT nuclei bypassed the microposts by forming deep local invaginations while preserving the overall nuclear shape.** Time-lapse movie of MEF WT nuclei stained with NucSpot Live 650 (a live-nuclear imaging dye, white) as they deform against rhodamine-labeled microposts (red) (Scale bar is 5 μm).

Movie 6: **WT deformation behavior is rescued upon expression of GFP lamin A in *Lmna^-/-^* cells.** Time-lapse movie of an MEF *Lmna^-/-^* nucleus expressing GFP-Lamin A (green) deforming against a rhodamine-fibronectin labeled micropost (red) and moving past it (corresponding to Figure 6C, third panel) (Scale bar is 10 μm).

Movie 7: **GFP-Lamin A (S22A/S392A) expression in *Lmna^-/-^* cells rescued WT deformation behavior but did not prevent entanglement.** Time-lapse movie of a MEF *Lmna^-/-^* nucleus (corresponding to Figure 6C, fourth panel) transfected with GFP-Lamin A (S22A/S392A) (green) and stained with NucSpot Live 650 (for DNA, blue), deforming around two rhodamine-fibronectin labeled microposts (red); white arrowheads show nuclear blebbing followed by NE rupture, as suggested by a decrease in cross-sectional area of the nucleus (Scale bar is 10 μm).

Movie 8: **Oil drop with higher surface tension deforming with a narrow, local invagination around the indenting wire.** The movie shows the behavior of an oil drop (blue) in water (yellow) when deformed with a thin metal wire (*diameter* = 0.5 mm) (Scale bar is 5 mm).

Movie 9: **Indentation with a metal wire of the oil drop with a lower surface tension causes a more extreme shape distortion from the starting oval shape.** Time-lapse movie shows the behavior of an oil drop (yellow) with 3% (w/v) Triton X-100 in water when deformed with a thin metal wire (*diameter* = 0.5 mm) (Scale bar is 5 mm).

## Acknowledgments

Microfabrication was performed with the assistance of the Nanoscale Research Facility at the University of Florida, and SEM imaging was performed at the Electron Microscopy Core, Interdisciplinary Center for Biotechnology Research at the University of Florida, and Microscopy and Imaging Center at Texas A&M University. A.K. thanks Brendan Mckee and Tanaya Roy for their help with cell culture, and Siddhika Chunchuwar, Ting-Ching Wang, and Dr. Qiao Zhang for their help with microscopy. This work was supported by NIH U01 CA225566 (T.P.L. and R.B.D.), a CPRIT established investigator award grant # RR200043 (T.P.L.), NIH R01 GM131178 (C.A.R.-K.), W.M. Keck Foundation award (C.A.R.-K.), R35GM126949 (K.J.R.), and NIGMS R01-GM123085 (P.P.L.), R.B.D.’s contribution is also based on work supported by (while serving at) the National Science Foundation.

## Data Availability Statement

The data that support the findings of this study are available from the corresponding authors upon reasonable request.

