## Supporting Data for "The nucleus bypasses obstacles by deforming like a drop with surface tension mediated by lamin A/C"

### Supporting information

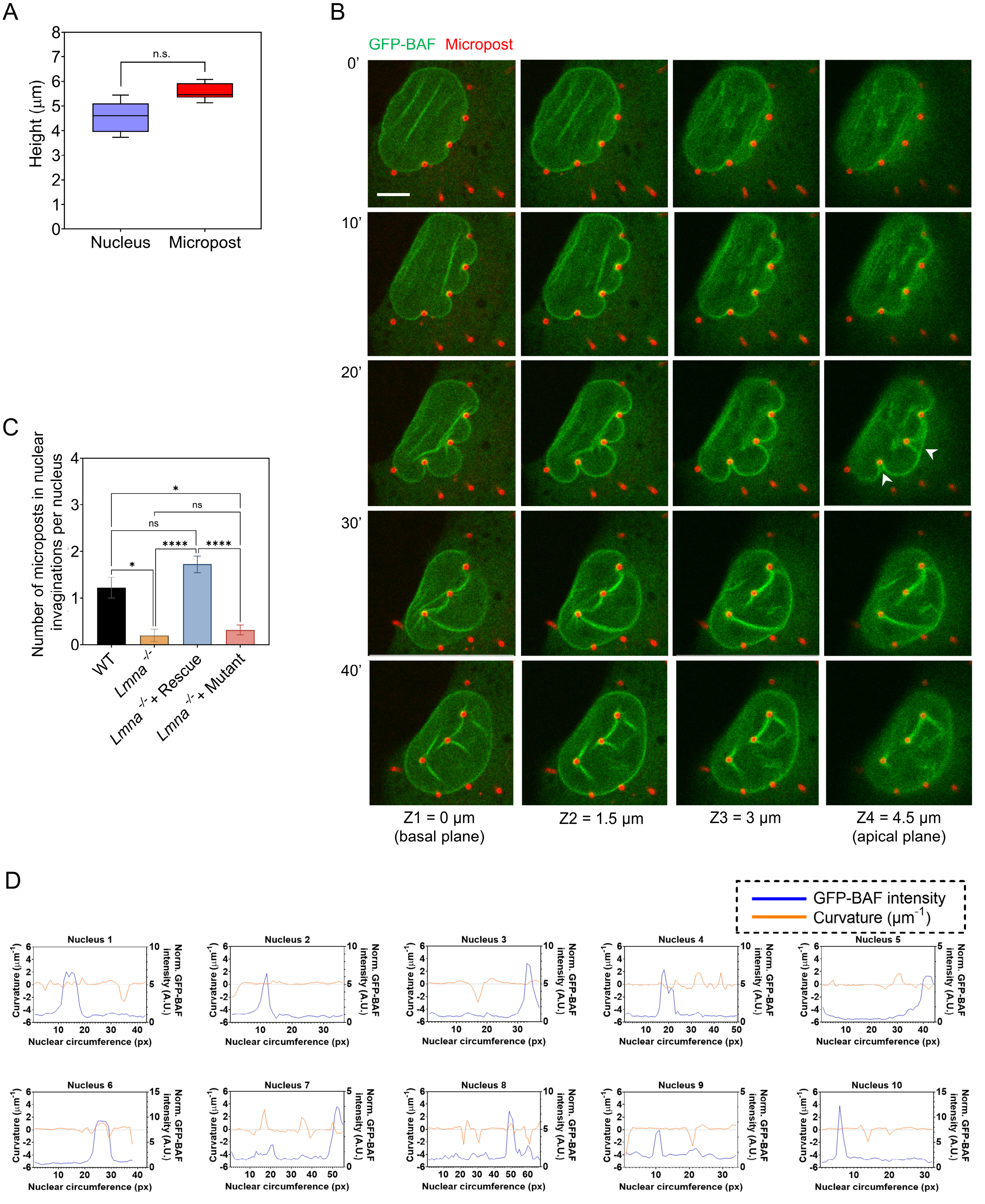


**Figure S1:** (A) Comparison of nuclear height measured in nuclei deformed around individual microposts with the height of microposts measured from reconstructed z-stacks; n ≥ 20 nuclei and n ≥ 20 microposts from at least three independent experiments. (ns: p > 0.05, Student's *t*-

test). (B) Time-lapse confocal images of an NIH 3T3 fibroblast stably expressing GFP-BAF deforming around 5 µm tall rhodamine-fibronectin stained PDMS microposts (red) sliding over the top of the microposts (white arrowheads), without collapsing the vertical microposts, (Scale bar is 5 µm). (C) Box plot shows the number of microposts in nuclear invaginations by the nucleus per cell in MEF WT (n = 12 cells), MEF *Lmna*$^{-/-}$ (n = 18 cells), MEF *Lmna*$^{-/-}$ + GFP-Lamin A (rescue) (n = 22 cells) and MEF *Lmna*$^{-/-}$ + GFP-Lamin A (S22A/S392A) mutant (n = 25 cells) from three experiments for each condition. (*: $p < 0.05$; Mann-Whitney test). (D) Plots of nuclear envelope curvature (orange curve) and GFP-BAF intensity (blue curve) along the circumference of nuclei (n = 10 cells) that deformed and underwent nuclear envelope rupture around microposts.

A

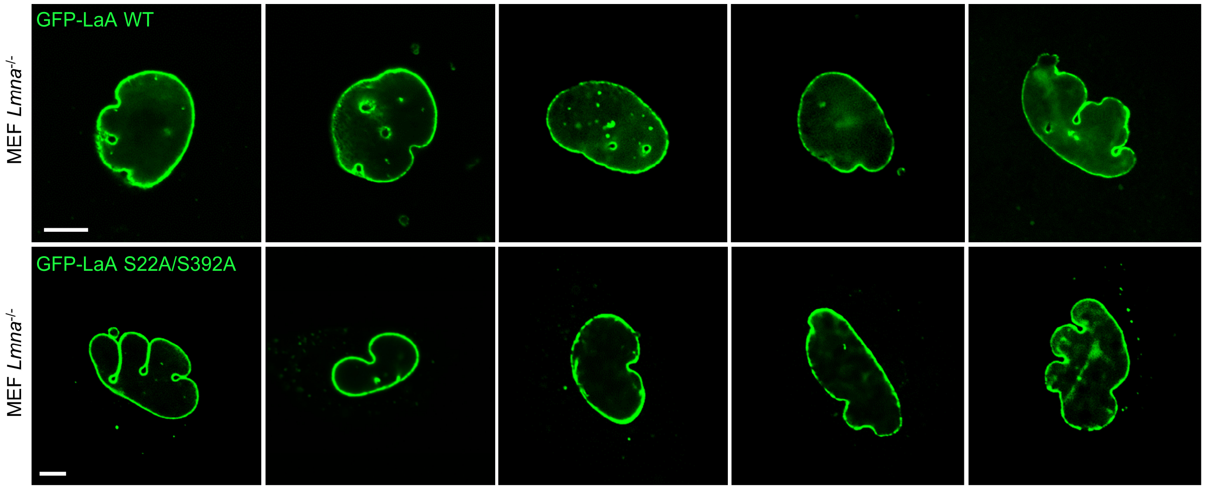


B

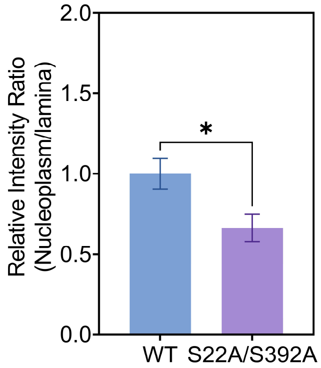


**Figure S2:** (A) Images of *Lmna*$^{-/-}$ nuclei expressing GFP-Lamin A WT in the top panel and GFP-Lamin A S22A/S392A in the bottom panel. (B) Bar graph compares nucleoplasmic to lamina intensity ratios of GFP-Lamin A (WT) and GFP-Lamin A S22A/S392A mutants (S22A/S392A), respectively. Data represents mean±SEM (n = 10 cells for each condition from three independent experiments, *: $p<0.05$, unpaired Student's *t*-test) normalized to the mean nucleoplasmic to lamina ratio of GFP-Lamin A.
